# Loss of the cytosolic DNA-sensing genes *CGAS* and *STING1* in armadillos (Cingulata)

**DOI:** 10.1101/2025.05.13.651073

**Authors:** Lillie Schaffer, Amanda Ivanoff, Frédéric Delsuc, Vincent J. Lynch

## Abstract

The principal sensor of intracellular double-stranded DNA (dsDNA) is cyclic GMP-AMP synthase (CGAS), which generates the second messenger cyclic GMP-AMP that binds stimulator of interferon genes (STING1), leading to the expression of type I interferon genes. CGAS and STING1 also play essential roles in maintaining genome integrity and the initiation and progression of cancer. Here we show that *CGAS* and *STING1* were pseudogenized in the ancestral armadillo branch 45 to 70 million years ago. The complete loss of the *CGAS*-*STING1* pathway in armadillos suggests this lineage has evolved alternate ways to sense intracellular double-stranded DNA, which may be related to their extreme cancer resistance.

## Introduction

Innate immunity is the first defense against foreign pathogens or endogenous danger signals, with distinct receptors for different danger signals, such as pathogen-associated molecular patterns (PAMPs) or damage-associated molecular patterns (DAMPs), that invoke an intracellular immune response. The principal sensor of intracellular double-stranded DNA (dsDNA) is cyclic GMP-AMP synthase (CGAS). CGAS binds to cytosolic dsDNA, generating the second messenger cyclic GMP-AMP (cGAMP), which then binds to the stimulator of interferon genes (STING1). cGAMP-bound STING subsequently undergoes a conformational change and translocates from the endoplasmic reticulum to the intermediate compartments between the endoplasmic reticulum and Golgi, which recruits TANK-binding kinase-1 (TBK1). TBK1 phosphorylates and activates interferon regulatory factor-3 (IRF3), leading to the expression of type I interferon genes^1^.

## Results

We recently completed a survey of gene losses in *Atlantogenata*^2^ and serendipitously noted that TOGA-based^3^ annotations indicated that the *CGAS* gene was lost in the Southern three-banded armadillo (*Tolypeutes matacus*), Mexican long-nosed armadillo (*Dasypus mexicanus;* formerly *D. novemcinctus*^4^), and giant anteater (*Myrmecophaga tridactyla)*, while *STING1* was lost in the Mexican long-nosed armadillo. To confirm these surprising observations, we used miniprot^5^ and minimap2^6^ to respectively map the Linnaeus’s two-toed sloth (*Choloepus didactylus*) CGAS and STING1 proteins and nucleotide coding sequences to 22 genome assemblies from 15 xenarthran species representing all major lineages (**Figure 1**). Unlike the TOGA-based annotation, the *CGAS* and *STING1* genes were complete and did not include inactivating mutations in sloths and anteaters (Pilosa). In contrast, *CGAS* had many inactivating mutations in six armadillo species (Cingulata), indicating it was pseudogenized and was completely missing in the chromosome-scale assembly of the Mexican long-nosed armadillo (*D. mexicanus*). *STING1* also had numerous inactivating mutations, including some shared by all seven armadillo species investigated. These data indicate that *CGAS* and *STING1* genes were pseudogenized in the ancestral armadillo branch 45 to 70 million years ago (Mya) (**Figure 1**). The complete loss of the CGAS-STING1 pathway in a major placental order (Cingulata) is only paralleled by pangolins (Pholidota) in mammals^7^.

**Figure 1:**
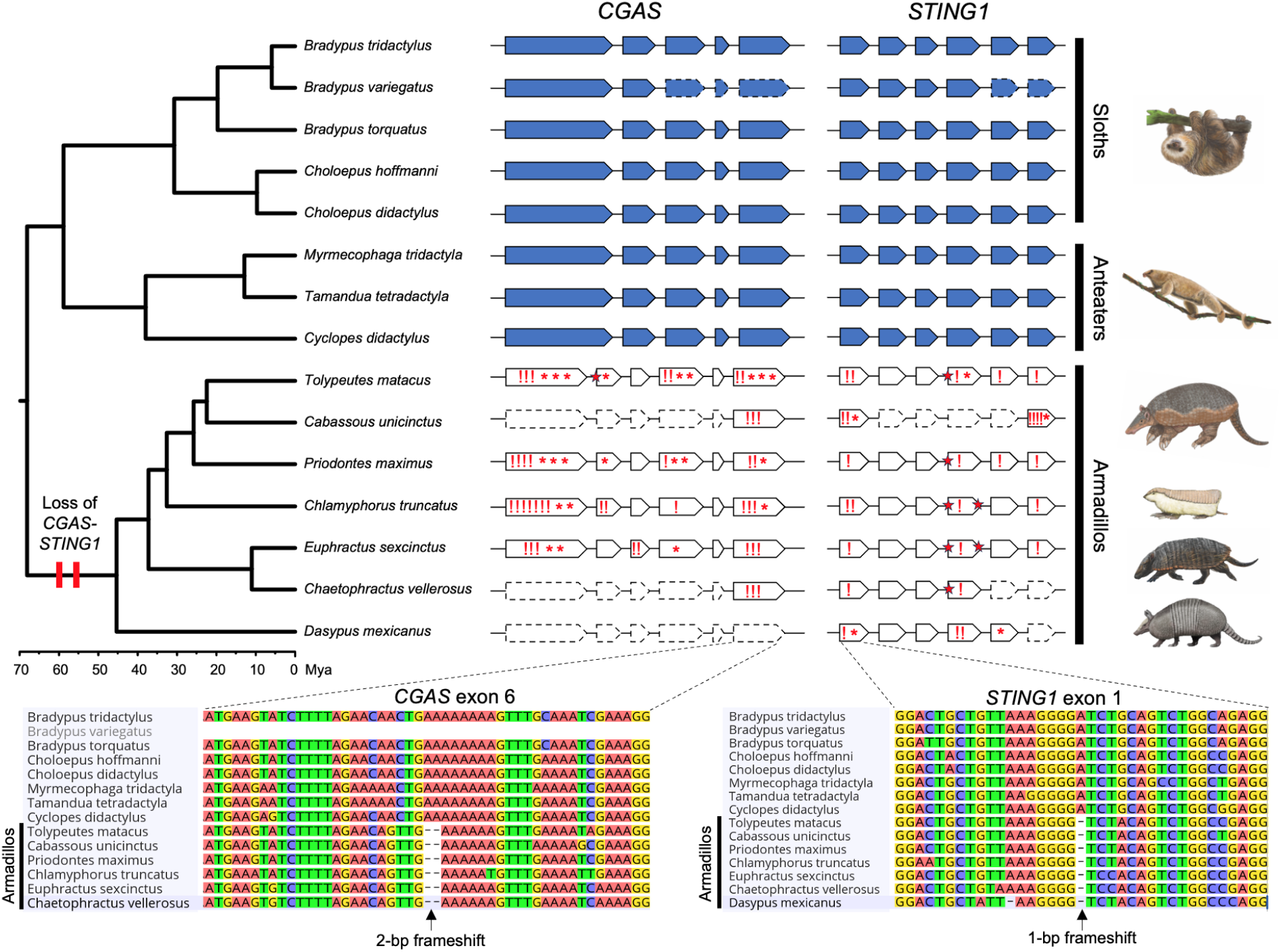
Evolutionary loss of *CGAS-STING1* in armadillos. Genomic organization of *CGAS* and *STING1* exons represented in the phylogenetic context of the 15 investigated xenarthran species (five sloths, three anteaters, and seven armadillos). Functional genes found in sloths and anteaters are represented in blue, and armadillo pseudogenes in white. Dashed line contours show exons missing in current genomic assemblies. Frameshifts (!), stop-codons (*), and splice site mutations (red stars) are indicated in armadillo pseudogenes. Sequence alignments present examples of inactivating mutations in exons shared by all armadillos, indicating that both *CGAS* and *STING1* were lost in their common ancestor 45 to 70 million years ago (Mya). Xenarthran phylogeny and timescale according to Gibb et al. (2016). Paintings by Carl Buell (copyright John Gatesy) and Michelle S. Fabros.

## Discussion

CGAS and STING1, widely recognized for their roles in the innate immune system^1^, also play essential roles in maintaining genome integrity and the initiation and progression of cancer. In the nucleus, for example, CGAS binds to chromatin^8–10^ and is recruited to sites of DNA double-strand breaks, where it inhibits RAD51-coated ssDNA filaments from initiating strand invasion^11^ and binds to PARP1 via poly(ADP-ribose), disrupting the formation of the PARP1–Timeless complex^12^. As a result of these processes, homologous recombination is suppressed, leading to increased chromosomal instability (CIN) and promoting tumorigenesis^11,12^. CIN also drives cancer metastasis^13–16^. Remarkably, chronic activation of the CGAS-STING1 pathway due to CIN rewires signaling in cancer cells, creating a pro-metastatic tumor microenvironment^17^. This rewiring causes rapid, diminishing responses (tachyphylaxis) to type-I interferon downstream of STING1; tumor cell-intrinsic STING1 inhibitors reduce CIN-driven metastasis in melanoma, breast, and colorectal cancers^17^.

CGAS and STING1 also play a central role in inflammation and senescence, in which cells with endogenous genomic DNA (gDNA) damage or mitochondrial (mt) dysfunction lead to the release of gDNA or mtDNA into the cytosol, causing activation of the CGAS-STING1 pathway. This can lead to permanent cell cycle arrest (senescence) and the release of inflammatory cytokines, also called the senescence-associated secretory phenotype (SASP)^18^. Prolonged activation of this pathway also leads to chronic systemic inflammation. For example, activation of the CGAS-STING pathway drives aging-related inflammation, neurodegeneration, and cognitive decline^18^. The absence of the CGAS-STING1 cytosolic pathway in armadillos and pangolins suggests that they may have evolved CGAS-STING1-independent mechanisms to respond to intracellular dsDNA, which may contribute to cancer resistance because the cancer-promoting effects of CGAS-STING have also been lost.

### Cancer Prevalence in Armadillos and Pangolins

While armadillos and pangolins do get cancer, it is remarkably rare in armadillos^19^. We compiled previously published data that systematically explore cancer prevalence in armadillos and found only two cases of neoplasia out of 342 necropsy reports, suggesting cancer prevalence is only around 0.58% (Clopper-Pearson exact 95% CI=0.071%–2.09%)^19–23^. An exhaustive literature search identified only a handful of cases of cancer in armadillos, including metastatic squamous cell carcinoma^24^, fibroma^25^, osteosarcoma^26^, adenocarcinoma in the stomach, mammary neoplasia with pulmonary metastasis^27^, a potentially virus-induced primary gastric T-cell lymphoma^28^, bronchiolar carcinoma, and a leiomyoma of the stomach^21^. Cancer can also be experimentally induced in armadillos—thalidomide treatment induced a highly malignant, aggressive choriocarcinoma that metastasized to the liver, mesentery, visceral parietal peritoneum, and lungs; the metastatic tumors in the lungs formed numerous teratomas^29^. In a similarly exhaustive literature search in pangolins, we found sporadic cases of benign hyperplasias, including a hemangioma^30^, an endometrial hyperplasia^31^, bile duct and thyroid gland hyperplasia^32^, and two cases of hepatocellular carcinoma^33^. A larger study of 22 wild-collected Chinese pangolins identified six cases of squamous cell carcinoma, three papillomas, at least one of which had early malignant changes, one case of adenomatoid, and several cases of hyperplasia of the gastric lining in the stomach. Unfortunately, there are no systematic studies in pangolins to reliably estimate their cancer prevalence. Thus, cancer is rare in armadillos, and possibly pangolins, which may be related to the loss of the evolutionary *CGAS* and *STING1* genes.

### Future Directions

The absence of the *CGAS* and *STING1* in armadillos and pangolins suggests that other genes in the pathway may also have been lost, or have altered patterns of molecular evolution, such as relaxed selective constraints or positive selection; characterizing patterns of molecular evolution in armadillos and pangolins might identify such shifts. It is also important to demonstrate that functions of CGAS and STING1 are not compensated by other genes; such inferences will require functional validation.

## Methods

### Inferences of Gene Loss Events

We downloaded TOGA loss_summ_data.tsv files, which used the human hg38 reference for each species, from https://genome.senckenberg.de/download/TOGA/. Gene losses in each species were identified as genes (GENE) annotated as “clearly lost” or “L” and which we coded as state 0, while all other TOGA categories were collapsed into a likely present category and coded as state 1; we note that this coding is conservative, as genes annotated with an “uncertain loss” and “partially intact” may be true losses. However, this coding scheme ensures we only infer losses with the highest confidence of true loss and may miss recent pseudogenization events when there are only one or a few inactivating mutations. Next, we inferred ancestral states and gene losses using the dollop program in Phylip (version 3.695), using the species phylogeny (with Paenungulata lineages as a polytomy) and enforcing all genes as present in the common ancestor; dollop infers ancestral dates using a Dollo’s law of irreversibility, in which once lost a gene cannot reevolve in descent lineages. We note that other processes could lead to an apparent reevolution of a previously lost gene such as incomplete lineage sorting, hybridization, or forms of gene flow such as horizontal gene transfer. While unlikely in mammals, Bayesian implementations of the Dollo model have been developed that account for processes like horizontal transfer^34,35^. To manually confirm these gene losses, we used miniprot^5^ and minimap2^6^ to map the Linnaeus’s two-toed sloth (*Choloepus didactylus*) *CGAS* and *STING1* proteins and nucleotide coding sequences to 22 genome assemblies from 15 xenarthran species representing all major lineages (Table S1). More specifically, miniprot was run with the following parameters: miniprot -j2 --aln --trans --gff to generate residue aligments, CDS predictions, and produce .gff annotation files detailing frameshifts, stop codons, and splice site mutations on predicted exons of each xenarthran assembly. Predicted CDSs were then aligned with MACSEv2^36^ and imported into GeniousPrime 2025.1.1^37^ to confirm frameshifts and stop codons. To further verify and visualize frameshifts and splice site mutations, minmap2 was run in Long-read spliced mode (-ax splice) to map CDSs to xenarthran assemblies within GeneiousPrime.

## Author Contributions (CRediT)

Conceptualization: V.J.L.

Data curation: L.S., A.N.I., F.D., V.J.L.

Formal analysis: L.S., F.D., V.J.L.

Funding acquisition: F.D., V.J.L.

Investigation: L.S., F.D., V.J.L.

Methodology: F.D., V.J.L.

Project administration: V.J.L.

Resources: L.S., F.D., V.J.L.

Software: N.A.

Supervision: V.J.L.

Validation: F.D., V.J.L.

Visualization: F.D.

Writing – original draft: F.D., V.J.L.

Writing – review & editing: L.S., A.N.I., F.D., V.J.L.

## Acknowledgments

This study was supported by NIH award 1R56AG071860-01 to VJL and grants from the European Research Council (ConvergeAnt project: ERC-2015-CoG-683257) and Investissements d’Avenir of the Agence Nationale de la Recherche (CEBA: ANR-10-LABX-25-01) to FD. This is contribution ISEM 2025-XXX of the Institut des Sciences de l’Evolution de Montpellier.

---

**Table S1:**
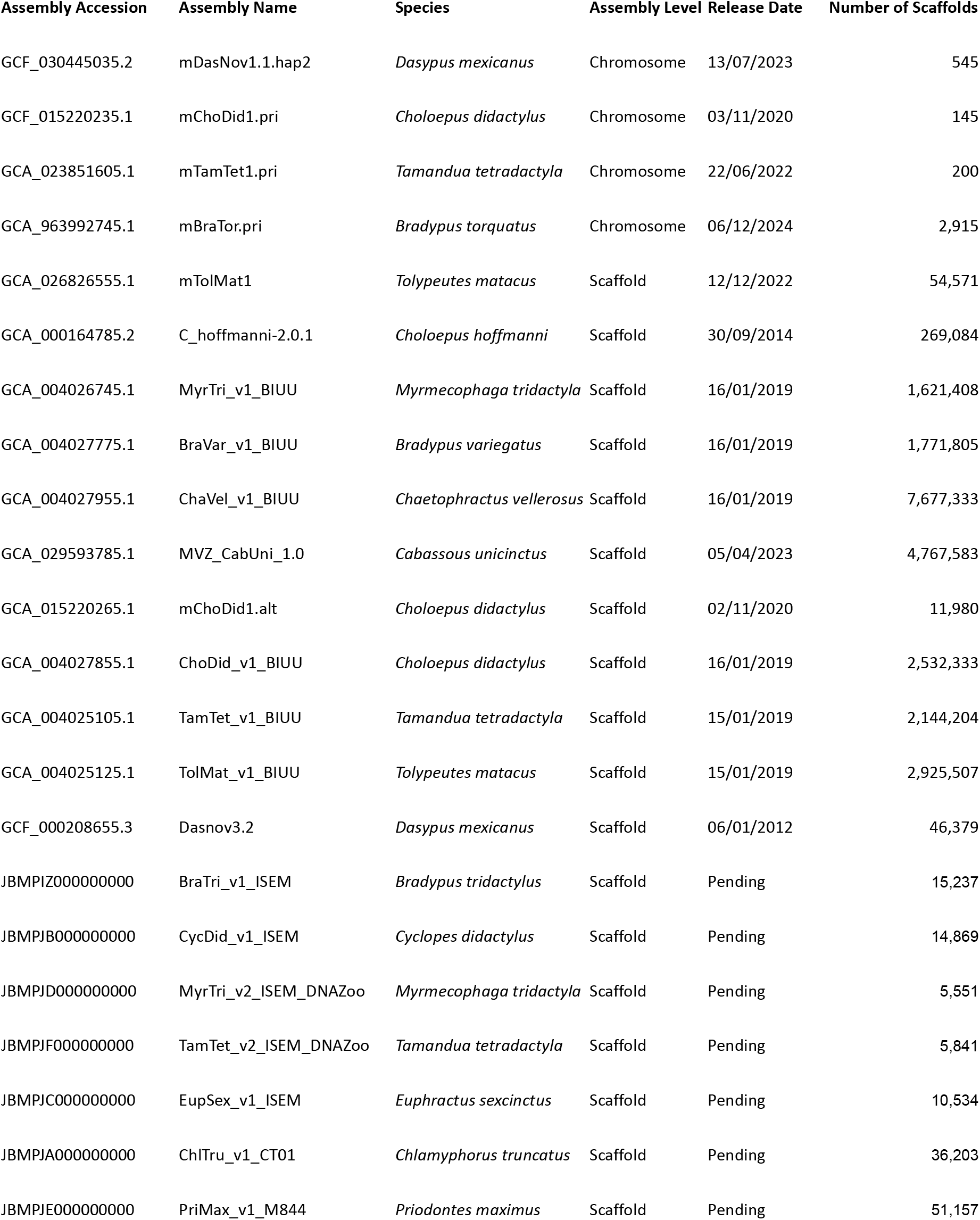
Details of xenarthran genome assemblies used.

| Assembly Accession | Assembly Name | Species | Assembly Level | Release Date | Number of Scaffolds |
| --- | --- | --- | --- | --- | --- |
| GCF_030445035.2 | mDasNov1.1.hap2 | *Dasypus mexicanus* | Chromosome | 13/07/2023 | 545 |
| GCF_015220235.1 | mChoDid1.pri | *Choloepus didactylus* | Chromosome | 03/11/2020 | 145 |
| GCA_023851605.1 | mTamTet1.pri | *Tamandua tetradactyla* | Chromosome | 22/06/2022 | 200 |
| GCA_963992745.1 | mBraTor.pri | *Bradypus torquatus* | Chromosome | 06/12/2024 | 2,915 |
| GCA_026826555.1 | mTolMat1 | *Tolypeutes matacus* | Scaffold | 12/12/2022 | 54,571 |
| GCA_000164785.2 | C_hoffmanni-2.0.1 | *Choloepus hoffmanni* | Scaffold | 30/09/2014 | 269,084 |
| GCA_004026745.1 | MyrTri_v1_BIUU | *Myrmecophaga tridactyla* | Scaffold | 16/01/2019 | 1,621,408 |
| GCA_004027775.1 | BraVar_v1_BIUU | *Bradypus variegatus* | Scaffold | 16/01/2019 | 1,771,805 |
| GCA_004027955.1 | ChaVel_v1_BIUU | *Chaetophractus vellerosus* | Scaffold | 16/01/2019 | 7,677,333 |
| GCA_029593785.1 | MVZ_CabUni_1.0 | *Cabassous unicinctus* | Scaffold | 05/04/2023 | 4,767,583 |
| GCA_015220265.1 | mChoDid1.alt | *Choloepus didactylus* | Scaffold | 02/11/2020 | 11,980 |
| GCA_004027855.1 | ChoDid_v1_BIUU | *Choloepus didactylus* | Scaffold | 16/01/2019 | 2,532,333 |
| GCA_004025105.1 | TamTet_v1_BIUU | *Tamandua tetradactyla* | Scaffold | 15/01/2019 | 2,144,204 |
| GCA_004025125.1 | TolMat_v1_BIUU | *Tolypeutes matacus* | Scaffold | 15/01/2019 | 2,925,507 |
| GCF_000208655.3 | Dasnov3.2 | *Dasypus mexicanus* | Scaffold | 06/01/2012 | 46,379 |
| JBMPIZ000000000 | BraTri_v1_ISEM | *Bradypus tridactylus* | Scaffold | Pending | 15,237 |
| JBMPJB000000000 | CycDid_v1_ISEM | *Cyclopes didactylus* | Scaffold | Pending | 14,869 |
| JBMPJD000000000 | MyrTri_v2_ISEM_DNAZoo | *Myrmecophaga tridactyla* | Scaffold | Pending | 5,551 |
| JBMPJF000000000 | TamTet_v2_ISEM_DNAZoo | *Tamandua tetradactyla* | Scaffold | Pending | 5,841 |
| JBMPJC000000000 | EupSex_v1_ISEM | *Euphractus sexcinctus* | Scaffold | Pending | 10,534 |
| JBMPJA000000000 | ChlTru_v1_CT01 | *Chlamyphorus truncatus* | Scaffold | Pending | 36,203 |
| JBMPJE000000000 | PriMax_v1_M844 | *Priodontes maximus* | Scaffold | Pending | 51,157 |

